# Auditory-motor entrainment and listening experience shape the perceptual learning of concurrent speech

**DOI:** 10.1101/2024.07.18.604167

**Authors:** Jessica MacLean, Jack Stirn, Gavin M. Bidelman

## Abstract

**Background:** Plasticity from auditory experience shapes the brain’s encoding and perception of sound. Though prior research demonstrates that neural entrainment (i.e., brain-to-acoustic synchronization) aids speech perception, how long- and short-term plasticity influence entrainment to concurrent speech has not been investigated. Here, we explored neural entrainment mechanisms and the interplay between short- and long-term neuroplasticity for rapid auditory perceptual learning of concurrent speech sounds in young, normal-hearing musicians and nonmusicians.

**Method:** Participants learned to identify double-vowel mixtures during ∼45 min training sessions with concurrent high-density EEG recordings. We examined the degree to which brain responses entrained to the speech-stimulus train (∼9 Hz) to investigate whether entrainment to speech prior to behavioral decision predicted task performance. Source and directed functional connectivity analyses of the EEG probed whether behavior was driven by group differences auditory-motor coupling.

**Results:** Both musicians and nonmusicians showed rapid perceptual learning in accuracy with training. Interestingly, listeners’ neural entrainment strength prior to target speech mixtures predicted behavioral identification performance; stronger neural synchronization was observed preceding incorrect compared to correct trial responses. We also found stark hemispheric biases in auditory-motor coupling during speech entrainment, with greater auditory-motor connectivity in the right compared to left hemisphere for musicians (R>L) but not in nonmusicians (R=L).

**Conclusions:** Our findings confirm stronger neuroacoustic synchronization and auditory-motor coupling during speech processing in musicians. Stronger neural entrainment to rapid stimulus trains preceding incorrect behavioral responses supports the notion that alpha-band (∼10 Hz) arousal/suppression in brain activity is an important modulator of trial-by-trial success in perceptual processing.

## INTRODUCTION

Everyday listening involves complex auditory scenarios in which listeners must isolate information from one talker in the presence of other talkers and background noise. Though difficult, many listeners successfully navigate these types of “cocktail party” listening environments. In particular, an extensive body of literature demonstrates perceptual advantages in speech-in-noise and “cocktail party” listening among highly-trained musicians (Bidelman & Yoo, 2020; Maillard et al., 2023; Parbery-Clark et al., 2009; Puschmann et al., 2018; Zendel & Alain, 2009). Despite evidence for a musician speech-in-noise advantage, the exact mechanism(s) underlying these enhancements are still under investigation. Enhanced sensory processing (Bidelman et al., 2011; Koelsch et al., 1999; Strait et al., 2010), attention (Román-Caballero et al., 2020; Strait & Kraus, 2011), and working memory/executive function (Kraus et al., 2012; Pallesen et al., 2010; Zuk et al., 2014) all might explain musicians’ superior figure-ground speech perception abilities.

One potential facilitatory mechanism that might enhance auditory perception, including noise-degraded and concurrent speech perception, is *neural entrainment*. Neural entrainment, or the yoking of ongoing neural oscillations to external stimuli, plays a strong role in governing the perceptual parsing of speech (Vanthornhout et al., 2018) and musical sounds (Doelling et al., 2019). Entraining to speech facilitates its intelligibility both in quiet and noise (Riecke et al., 2018). On the contrary, electrophysiological studies have shown that poorer entrainment in clinical populations (e.g., listeners with auditory processing disorder) parallels behavioral deficits in concurrent speech listening tasks (Gilley et al., 2016; Momtaz et al., 2021; Momtaz et al., 2022). Collectively, these studies suggest the robustness of the brain’s neuroacoustic entrainment might play an important role in successfully parsing concurrent speech signals.

In addition to their improved speech-in-noise performance, studies demonstrate musicians have stronger neural entrainment to musical stimuli in the beta range (∼20 Hz) that are associated with rhythmic processing and forming temporal predictions (Doelling & Poeppel, 2015). Such rhythmic brain oscillations have also been linked to sensorimotor synchronization ability (Arnal et al., 2014; Krause, Pollok, et al., 2010; Krause, Schnitzler, et al., 2010). Indeed, a relationship between enhanced neural entrainment strength and successful sensorimotor synchronization (such as tapping along with a beat) has also been demonstrated on an individual level (Nozaradan et al., 2016). In addition to the beta band, modulations in alpha entrainment (∼10 Hz) can also influence speech perception. Increases in alpha brain rhythms are traditionally associated with internal reflection or decreased attention to a given task (Klimesch, 2012). Related to task performance, phase-related changes and *suppression* of alpha activity have been shown to predict successful speech intelligibility in quiet and noise (Obleser et al., 2012; Strauß et al., 2015; Weisz et al., 2011). Germane to our current study, Puschmann and colleagues found that when attending to continuous speech in quiet, the amount of participants’ music training positively correlated with the strength of alpha-band phase locking between the primary auditory cortex and dorsal and ventral auditory pathways, suggesting alpha-band entrainment to speech across the cortex is influenced by music training (Puschmann et al., 2021). Cortical alpha states also influence brainstem speech encoding through dynamic fluctuations in arousal and attention, and consequently, are relevant to speech processing at multiple stages of the auditory system (Lai et al., 2022). Thus, as evidenced by changes in several prominent time-frequency signatures of the EEG, studies support the notion that musicianship might enhance the brain’s ability to entrain to external acoustic sounds. Consequently, the current study aimed to determine whether musicians’ extensive experience with sensorimotor synchronization and enhanced neural entrainment might also enhance aspects of concurrent speech listening.

In addition to speech-to-brain entrainment, brain-to-brain interactions between the auditory and motor systems might also aid the perception of “cocktail party” speech. Previous studies have demonstrated engagement of the motor system (alongside the auditory system) to enhance the neural representation of speech (Poeppel & Assaneo, 2020; Poeppel & Hickok, 2004). Indeed, close coordination between the premotor and temporal cortices is used to track various linguistic elements of the speech signal spanning the syllable, word, and phrase levels (Assaneo & Poeppel, 2018; Ding et al., 2016; He et al., 2023; Keitel et al., 2018). Motor engagement is particularly evident under noise degradation when efference copy must enhance speech representations from the impoverished acoustic input (Du et al., 2014). Such top-down, cross-modal enhancement of auditory information might also be due to the ability of the motor system to enhance temporal predictions of sensory stimuli (Dick et al., 2011; Morillon & Baillet, 2017). These mechanisms could presumably improve degraded listening skills. One idea is that the enhanced auditory-motor integration necessary for musicians may enhance auditory-motor connectivity, thus enabling their more successful speech-in-noise comprehension (e.g., Du & Zatorre, 2017).

Functional connectivity between the auditory and motor systems (i.e., the degree of coupling between regional activity) can be used to directly characterize auditory-motor signaling. Indeed, connectivity within the alpha band that tags the speech signal is stronger in musically trained individuals (Puschmann et al., 2021). Functional connectivity enhancements gained through long-term music training may even align with prevention of typical age-related declines in speech-in-noise perception (Zhang et al., 2024). In addition to connectivity strength, the direction of signaling (i.e., auditory-to-motor vs. motor-to-auditory) can provide insight into “bottom up” vs. “top-down” mechanisms of auditory-motor involvement. Stronger connectivity in the auditory-to-motor direction could indicate greater reliance on sensory cue extraction and specific stimulus features, whereas stronger motor-to-auditory signaling could indicate greater reliance on predictive or anticipatory cues to perceive cocktail party speech.

In the present study, we reanalyzed the EEG data collected in our previously published study on the neuroplasticity of concurrent speech sound learning in musicians and nonmusicians (MacLean et al., 2024). In our prior work, we found that long-term plasticity (e.g., musicianship) interacted with short-term perceptual learning (e.g., learning a task within one ∼45 minute session) in the perception of double-vowel speech stimuli. Musicians and nonmusicians demonstrated different cortical (but not subcortical) learning trajectories which related to behavioral measures of speech identification success. Fortuitously, our stimulus design included a rapid cueing speech train which had the natural potential to induce neural entrainment prior to listeners’ behavioral decision (Bidelman, 2015). Here, we performed novel analyses on this critical segment to understand how neural entrainment and auditory-motor connectivity interact with long-and short-term auditory experiences during double-vowel learning. We hypothesized that alpha-band entrainment to concurrent speech and auditory-motor connectivity strength would influence behavioral success at the single trial level, and that these oscillatory processes would differ between musician and nonmusician groups. Additionally, we investigated the direction of auditory-motor connectivity to understand how the relative signaling between the auditory and motor systems relate to learning and behavioral performance. Our findings support the notion that alpha suppression is critical for task success and bottom-up use of stimulus features for both groups, while revealing differences in neural entrainment based on long-term music training.

## MATERIALS AND METHODS

The current study represents a new analysis of neural entrainment from the EEG data reported in MacLean et al. (2024). Evoked potential results including brainstem (FFR) and cortical (ERP) responses to speech and how they are modulated by perceptual learning are reported in the companion paper (MacLean et al., 2024). The reader is referred to the original manuscript for full methodological details.

### Participants

Twenty-seven young adults (ages 18-34; mean ± *SD*: 23.68 ± 4.22; 13 female) with normal hearing thresholds (bilateral pure tone averages < 25 dB HL, octave frequencies between 250 and 8000 Hz) participated in this study. All participants were fluent in American English and reported no previous neurologic or psychiatric disorders. Participants gave written, informed consent in accordance with a protocol approved by the Indiana University Institutional Review Board.

Participants were separated into musician (M; *n* = 13) and nonmusician (NM; *n* = 14) groups based on their extent of formal music training. Musicians had at least 10 years of formal music training starting at or before age 12, while nonmusicians had 5 or fewer years of lifetime music training (Wong et al., 2007). Groups did significantly differ in amount of music training (M: 16.1 ± 4.3 years; NM: 2.4 ±1.7 years; *t*(25) = 10.93; *p* < 0.001), but were matched in age (*t*(25) = 1.58; *p* = 0.413), cognitive ability as assessed through the Montreal Cognitive Assessment (Nasreddine et al., 2005) (*t*(25) = 1.78; *p* = 0.088), self-reported bilingualism (*X^2^*(1, *N* = 27) = 0.022, *p* = 0.883), sex balance (*X ^2^*(1, *N* = 27) = 1.78, *p* = 0.182), and handedness as assessed through the Edinburgh Handedness Inventory (*t*(25) = -0.615; *p* = 0.544) (Oldfield, 1971).

### Double-vowel stimuli and task

Concurrent vowel stimuli were modeled after previous studies (Alain et al., 2007; Assmann & Summerfield, 1989, 1990; Bidelman & Yellamsetty, 2017). Stimuli consisted of synthesized, steady-state vowels (/a/, /e/, and /i/) which were presented in three unique vowel combinations (i.e., /a/ + /e/; /e/ + /i/; /a/ + /i/). Vowels were never paired with themselves. Stimuli were created with a Klatt-based synthesizer (Klatt, 1980) coded in MATLAB (v 2021; The MathWorks, Inc., Natick, MA). Each vowel was 100 ms in duration with 10-ms cos^2^ onset/offsetramping to prevent spectral splatter. The fundamental frequency (F0) between vowels was 4 semitones (150 and 190 Hz), which promotes segregation for most listeners (Assmann & Summerfield, 1990; Bidelman & Yellamsetty, 2017). F0 and the first two formant frequencies (*F1*a,e,i = 787, 583, 300 Hz; *F2* a,e,i = 1307, 1753, 2805 Hz) remained constant for the duration of the token.

The speech sounds were presented in rarefaction phase through a TDT RZ6 interface (Tucker-Davis Technologies, Alachua, FL) controlled via MATLAB. Stimuli were presented binaurally at 79 dB SPL through electromagnetically shielded (Campbell et al., 2012; Price & Bidelman, 2021) ER-2 insert earphones (Etymotic Research, Elk Grove, IL). Prior to EEG testing, we required all participants to identify single vowels with 100% accuracy. This ensured subsequent learning would be based on improvements in *concurrent* speech identification rather than isolated sound labeling ability.

We used a clustered stimulus paradigm (Bidelman, 2015) employing interspersed fast and slow interstimulus intervals (ISIs) to collect speech-evoked potentials during the active perceptual task (**Figure 1**). Speech-ERP/FFR data are reported in the companion paper (MacLean et al., 2024). Each trial consisted of one of the three vowel combinations. During a trial, 20 repetitions of the vowel pair were presented with a fast ISI of 10 ms to elicit the FFR. Thus, the corresponding stimulus onset asynchrony (SOA) was 110 ms (i.e., 9.09 Hz). The ISI was then slowed to 1100 ms and a single stimulus was presented to evoke the ERP and cue a behavioral response. Participants then identified both vowels through keyboard responses following the isolated vowel pair. The next trial began after the participants’ response and 250 ms of silence. Participants were asked to identify both vowels as quickly and accurately as possible (no feedback was provided). Double vowel pairs were randomized in order. This identical task was repeated over four learning blocks. In total, each block included 150 stimulus trials. Each block took 10-15 min to complete. Participants were offered a short (2-3 min) break after each block to avoid fatigue.

**Figure 1.**
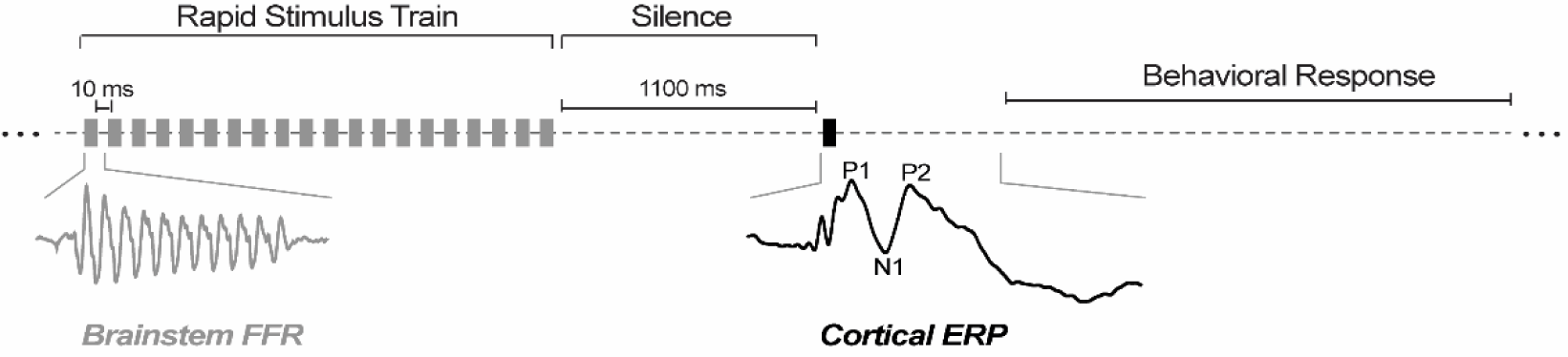
Clustered stimulus paradigm to induce alpha-band speech entrainment. The stimulus paradigm began with a rapid stimulus train presented at ∼10 Hz, followed by a 1100 ms period of silence before the isolated vowel pair which cued behavioral responses (Bidelman, 2015). Analyses were performed on induced neural entrainment observed during the silent period.

To investigate neural entrainment induced by the preceding speech stimuli prior to listeners’ behavioral response, we isolated neural activity to the silent stimulus portion (-1100 to 0 ms) immediately following the rapid speech train. This allowed us to assess how ongoing brain rhythms that have entrained to speech after its cessation modulate subsequent success in identification.

### EEG recording and preprocessing

We used Curry 9 (Compumedics Neuroscan, Charlotte, NC) and BESA Research 7.1 (BESA, GmbH) to record and preprocess the continuous EEG data. Continuous EEGs were acquired from 64-channel Ag/AgCl electrodes positioned at 10-10 scalp locations (Oostenveld & Praamstra, 2001). Recordings were digitized at 5 kHz using Neuroscan Synamps RT amplifiers. Data were referenced to an electrode placed 1 cm behind Cz during online recording. Data were re-referenced to common average reference for subsequent analysis. Impedances were kept below 25 kΩ. Electrodes placed on the outer canthi of the eyes and superior and inferior orbit captured ocular movements. Eyeblinks were corrected using a topographic principal component analysis (Wallstrom et al., 2004). Responses were collapsed across vowel pairs to obtain an adequate number of trials for analysis (Bidelman & Yellamsetty, 2017; Yellamsetty & Bidelman, 2018). Responses exceeding 150 µV were rejected as further artifacts. We then bandpass filtered responses from 7 to 12 Hz (zero-phase Butterworth filters; slope = 48 dB/octave) to isolate alpha-band activity (Alain et al., 2023; Bidelman, 2017; Lai et al., 2022), corresponding to the nominal rate of our speech train stimuli. Data were then epoched during the silent portion of the stimulus presentation (-1100 to 0 ms), baselined, and ensemble averaged to derive sustained response waveforms for each condition per subject. For subsequent analyses, neural responses were separated by listeners’ trial-by-trial response accuracy (correct vs. incorrect trials).

### Fast-Fourier Transforms

To measure the strength of neural entrainment induced by the rapid stimulus train, we computed the Fast Fourier Transform (FFT) in the -1100 to 0 ms time window separately for each block and correct/incorrect trials. We measured the magnitude and frequency for the maximum spectral peak within the alpha band (7-12 Hz) at the Cz electrode to quantify entrainment at the scalp level.

### Functional connectivity

To resolve the underlying brain sources of entrainment effects, we measured directional flow of information within auditory-motor networks using Granger Causality (GC) (Geweke, 1982; Granger, 1969). GC measures the degree to which Signal A “Granger-causes” Signal B and is computed directionally in order to infer causal flow of information between brain regions. We computed functional connectivity in the frequency domain between primary auditory (A1) and motor (M1) cortices, bilaterally, using BESA Connectivity (v2.0) (Dhamala et al., 2008; Geweke, 1982). A1 and M1 regions of interest (ROI) were defined via Talairach coordinates in template brain space (*x, y, z* coords.: M1: ±44.8, -7.8, 38.24 cm; A1: ±50.4, -21.7, 11.5 cm). Frequency decomposition was based on complex demodulation (Papp & Ktonas, 1977), which results in uniform frequency resolution across the analysis bandwidth (i.e., sliding window FFT). The time-frequency analysis initially spanned the entire epoch window (-3400 to 1000 ms), using a pre-stimulus baseline (-3400 to -3200 ms) over a bandwidth between 5-20 Hz (i.e., centered at the nominal alpha frequency). However, we extracted GC within the post-stimulus train silence (-1100 to 0 ms) within the 7-12 Hz band (collapsed across time and frequency) to examine alpha auditory motor coupling just prior to the target cue and behavioral decision. We computed GC between A1 and M1 in both the forward and reverse directions (A1 → M1 and M1 → A1, respectively) to assess directed “bottom-up” and “top-down” neural signaling between auditory and motor system.

### Statistical analyses

Unless otherwise noted, we analyzed dependent variables using mixed model ANOVAs in R (version 4.2.2) (R-Core-Team, 2020) and lme4 package (Bates et al., 2015). Behavioral measures (percent correct, reaction time) were analyzed with fixed effects of group (2 levels), block, (4 levels), Entrainment strength was analyzed with fixed effects of group (2 levels), block (4 levels), behavioral response (2 levels, correct or incorrect), and random effect of subject. Additionally, we included a covariate for the number of trial counts for correct and incorrect responses. Granger Connectivity was analyzed with the same fixed and random effects as above, with two additional fixed effects of hemisphere (2 levels; left vs. right) and direction (2 levels; forward: A1 → M1, reverse: M1 → A1). Effect sizes are reported as partial eta squared (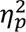) and degrees of freedom (*d.f.*) using Satterthwaite’s method. Multiple pairwise comparisons were adjusted using Tukey method. Linear contrasts were adjusted using the Sidak method.

Initial diagnostics indicated heavy tailed distributions for both neural measures. Consequently, we used the Box-Cox procedure (Box & Cox, 1964) to transform the data and satisfy normality assumptions necessary for parametric statistics. This procedure transforms the data according to y’=(y^λ^ – 1)/λ, where λ = 0.071 and λ = -0.11 where determined empirically for entrainment strength and connectivity, respectively.

## RESULTS

### Behavior

**Figure 2** displays behavioral results for both M and NM groups across all four training blocks. An ANOVA on behavioral accuracy revealed a main effect of training block [*F*(3, 75) = 12.13, *p* < 0.001, 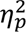= 0.33], where both groups improved in accuracy with training block (linear contrast: M: *t*(75) = 4.34, *p* < 0.001; NM: *t*(75) = 3.01, *p* = 0.0035). These data demonstrate that both groups improved in speech identification accuracy with training.

**Figure 2.**
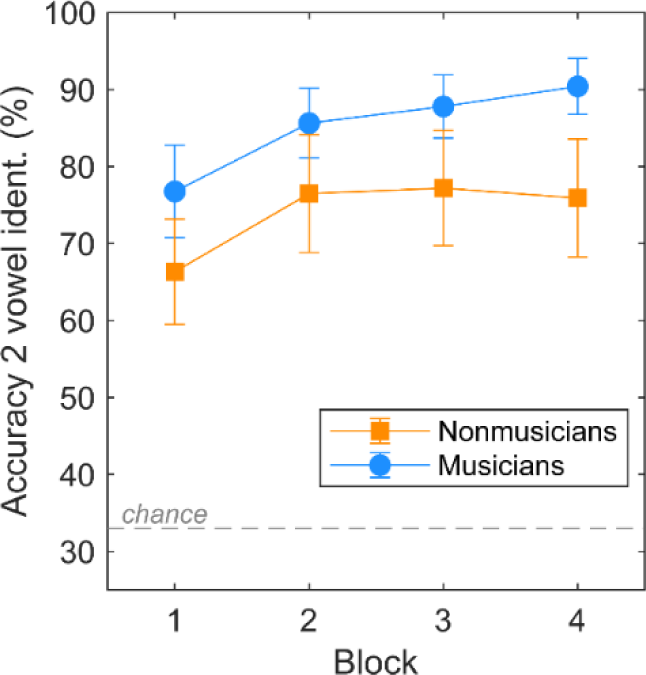
Behavioral accuracy increased with training block. Accuracy in identifying both concurrently presented vowels improved with increasing training block for musicians and nonmusicians. Error bars = ±1 S.E.M. Reprinted from MacLean et al. (2024), with permission from Oxford University Press.

### Neural entrainment to speech preceding behavior

**Figure 3** displays alpha-band waveforms during the entire stimulus period. For subsequent analyses, we focused on the silent period just prior to the target double-vowel presentation that cued listeners’ behavioral response.

**Figure 3.**
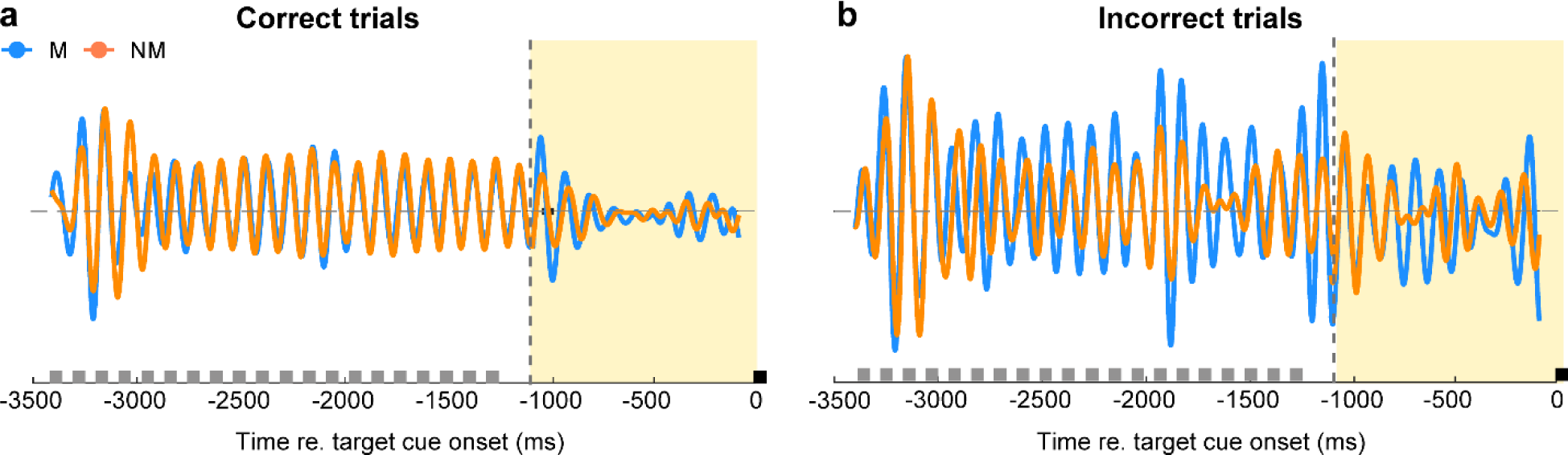
Alpha-band entrainment during the stimulus time course. Alpha-band (7-12 Hz) entrainment waveforms for musicians and nonmusicians preceding correct (**a**) and incorrect (**b**) behavioral responses. Gray boxes represent stimulus in the rapid stimulus train (see Fig. 1). To assess true entrainment, analyses were performed during the silent portion of the stimulus paradigm (yellow) just prior to the behavior-cueing token at *t* = 0 (black box).

**Figure 4** shows alpha-band entrainment amplitude in the pre-stimulus silence period (i.e., just prior to the target cue) for correct and incorrect trials. An ANOVA on alpha-band entrainment amplitude revealed a 2-way interaction between group x trial accuracy [*F*(1, 172.38) = 4.52, *p* = 0.035, 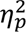= 0.03] (**Fig. 4b**), driven by larger spectral amplitudes for musicians than nonmusicians preceding incorrect trials [*t*(29) = 2.20, *p* = 0.036]. Groups showed similar response amplitudes before correct trials [*t*(29) = 1.011, *p* = 0.32]. There was also a block x response accuracy interaction [*F*(3, 172.38) = 4.014, *p* = 0.0086, 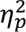 = 0.07]. A linear contrast revealed this interaction was due to a steady increase in response amplitude across blocks for incorrect trials [*t*(177) = 4.18, *p* < 0.0001], regardless of group. Responses were invariant across blocks for correct trials [*t*(177) = -0.41, *p* =0.68].

**Figure 4.**
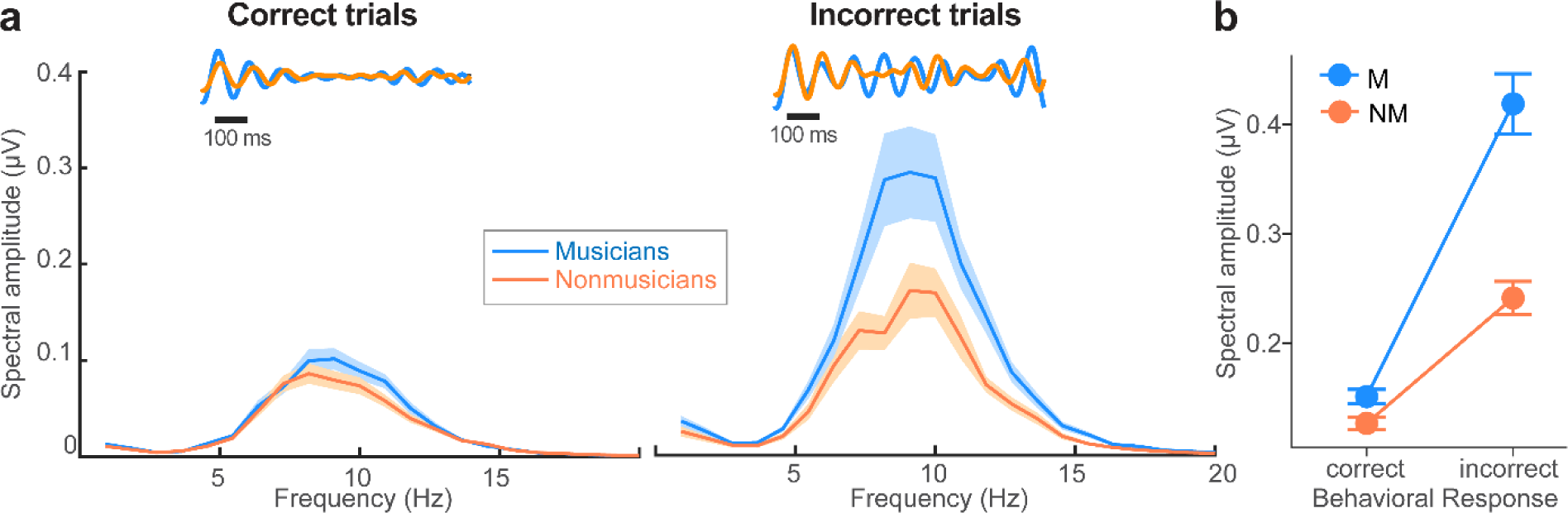
Neural entrainment following a rapid speech stimulus train predicts subsequent behavioral identification accuracy for double-vowel mixtures. (a) **FFTs are displayed for** musicians and nonmusicians for silences preceding correct and incorrect trials. Insets show time waveforms of the post-train period (see yellow shading, Fig. 3). **(b)** Musicians had stronger entrained responses preceding incorrect trials than did nonmusicians, despite similar responses preceding correct trials. Error bars/shading = ±1 S.E.M.

### Auditory-motor connectivity

**Figure 5** depicts time-frequency plots of source-level waveforms from auditory (A1) and motor (M1) cortex in the left (LH) and right hemispheres (RH) per group. Spectrographic maps were used to calculate Granger Connectivity (GC), reflecting directed neural signaling between ROIs, for correct and incorrect trials per hemisphere and group (**Figure 6**).

**Figure 5.**
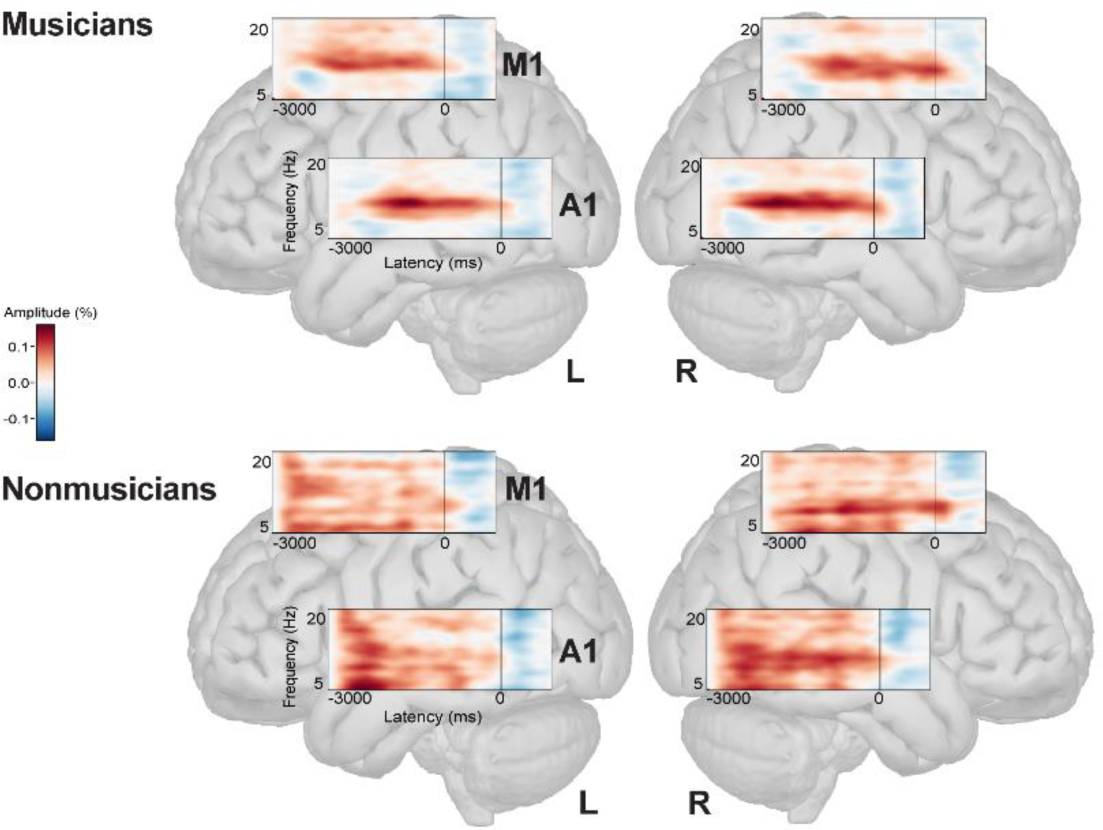
Source time-frequency responses reflecting neural entrainment within the auditory-motor network. Each spectrogram demonstrates spectral density within the alpha band range stemming from auditory (A1) and motor (M1) cortex. Hot colors, %-increase in activity relative to baseline; cool colors, %-decrease activity. *t* = 0 denotes the onset of the double-vowel mixture that cued listeners’ behavioral response. Note the power at ∼10 Hz reflecting phase-locking to the rapid stimulus train (see Fig. 1a) which is also stronger in musicians. L/R = left/right hemisphere.

**Figure 6.**
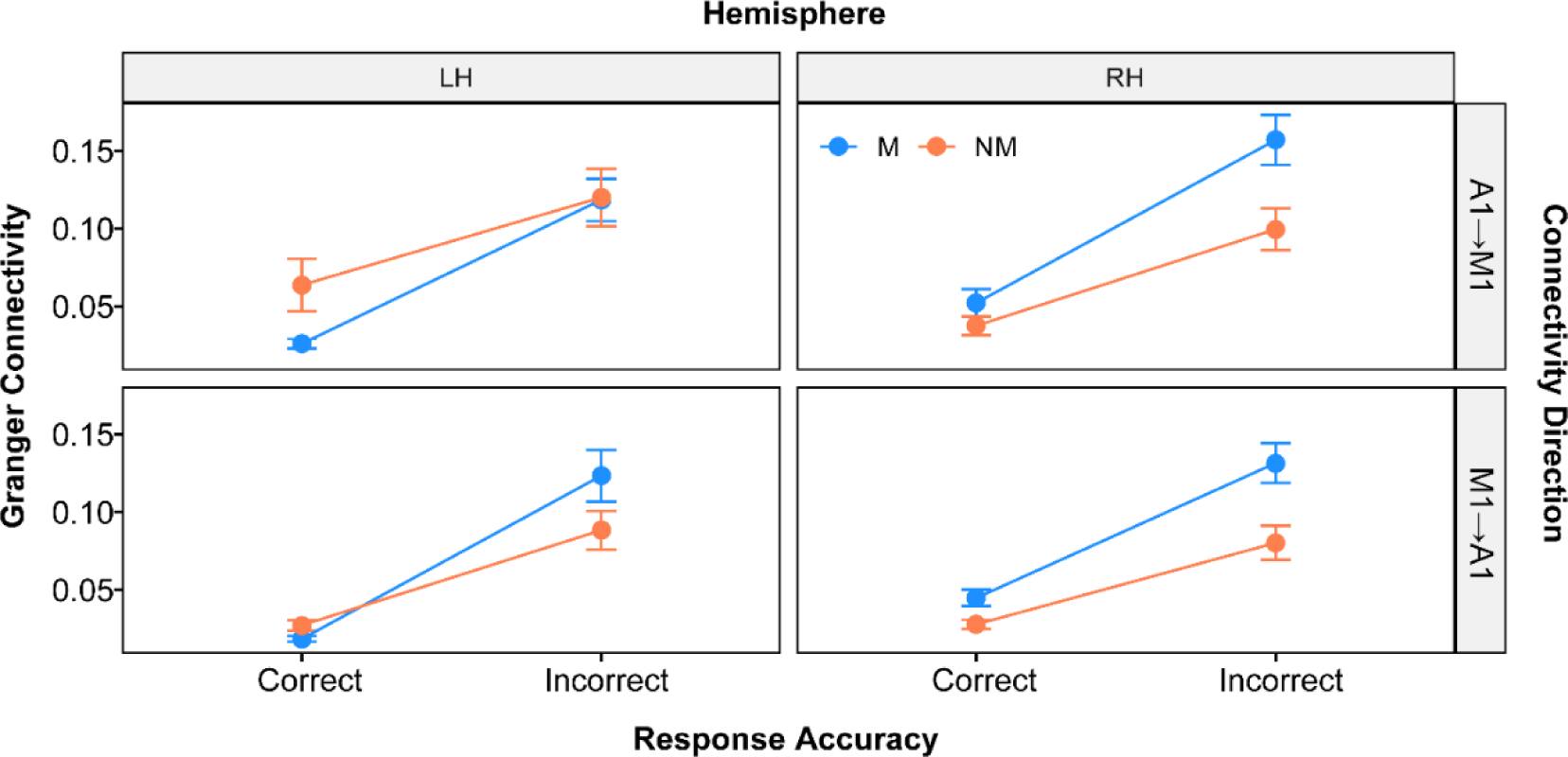
Auditory-motor coupling varies by group, hemisphere, and trial-wise accuracy. Granger connectivity values were strongest for musicians in the right hemisphere preceding incorrect trials. Both groups had weaker connectivity in the motor to auditory direction. Errorbars = ±1 S.E.M.

A mixed-model ANOVA on GC strength revealed several two-way interactions (**Figure 7**). We found an interaction between hemisphere and group (**Fig. 7a**) [*F*(1, 766.72) = 9.07, *p* = 0.0027, 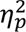= 0.01]. This was driven by stronger GC values in the right hemisphere compared to the left hemisphere for musicians only (pairwise comparison: M: *t*(766) = -4.69, *p* < 0.0001, NM: *t*(766) = - 0.555, *p* = 0.58). An interaction between group and block (**Fig. 7b**) [*F*(3, 769.72) = 3.64, *p* = 0.013, 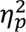 = 0.01] was driven by greater connectivity with block for musicians only (linear contrast: M: *t*(791) = 3.16, *p* = 0.0050; NM: *t*(779) = 1.51, *p* = 0.35). We also observed an interaction between group and response accuracy (**Fig. 7c**) [*F*(1, 766.72) = 5.030, *p* = 0.025, 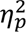 < 0.01] which was driven by stronger GC during incorrect trials in both groups but especially musicians (pairwise comparison: M: *t*(766) = -13.69, *p* < 0.0001; NM: *t*(766) = -11.11, *p* < 0.0001). Finally, an interaction between block and response accuracy (**Fig. 7d**) [*F*(3, 766.72) = 4.55, *p* = 0.0036, 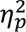= 0.02] was driven by increasing connectivity with block for incorrect trials only (linear contrast: correct: *t*(787) = 0.12, *p* = 0.99; incorrect: *t*(787) = 4.60, *p* < 0.0001). All other interactions were non-significant. Additionally, we observed a main effect of direction (**Fig. 7e**) [*F*(1, 766.72) = 4.53, *p* = 0.034, 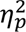 < 0.01], attributed to higher auditory-to-motor (i.e., A1 → M1) compared with motor-to-auditory (i.e., M1 → A1) connectivity in both groups.

**Figure 7.**
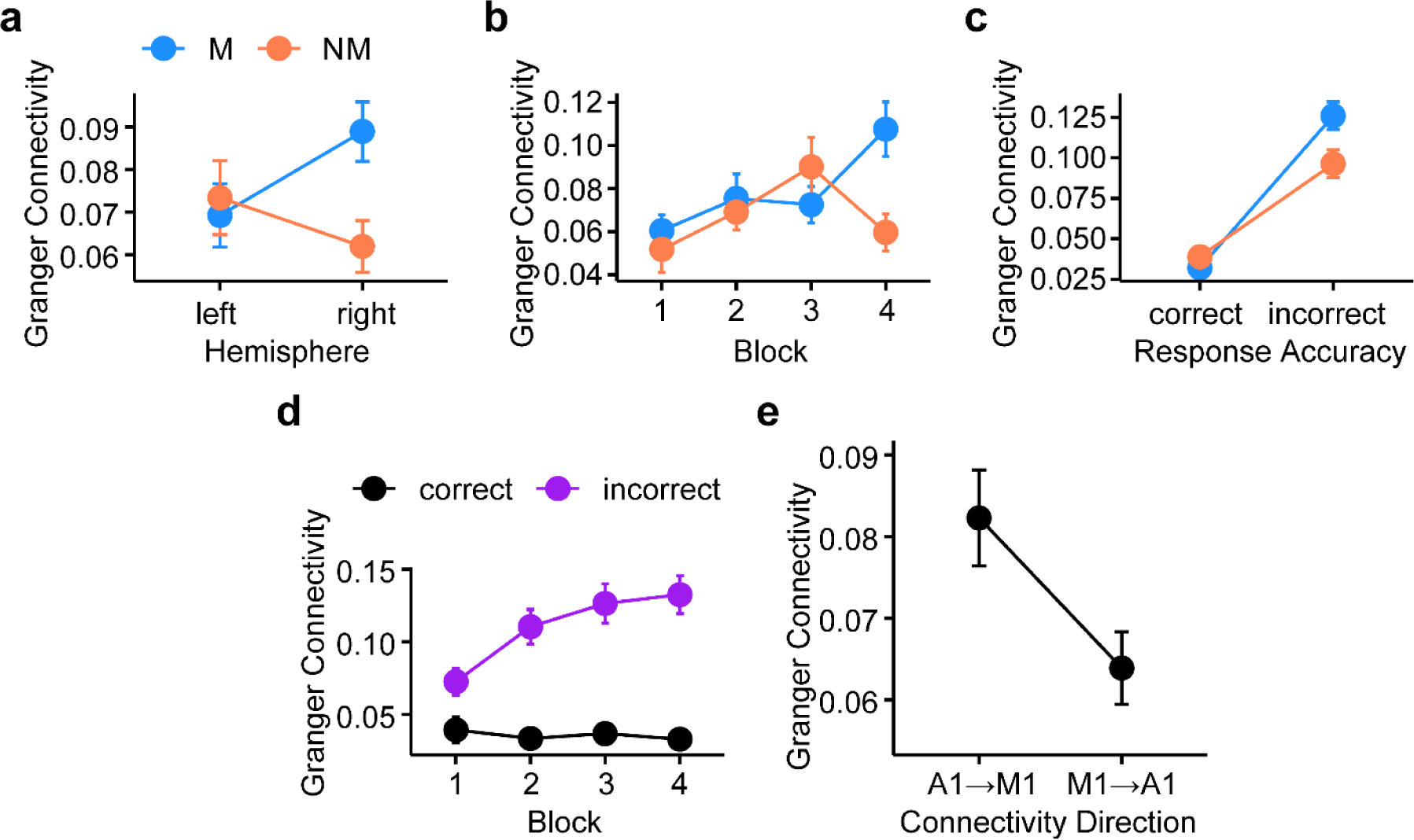
Significant interactions and main effects on Granger Connectivity. (**a**) Hemisphere x group, (**b**) block x group interaction, (**c**) group x response accuracy, and (**d**) block x response accuracy interactions. (**e**) Main effect of direction. Connectivity was stronger in the auditory-motor (bottom-up) vs. motor-auditory (top-down) direction. Errorbars = ±1 S.E.M.

## DISCUSSION

By analyzing EEG entrainment during perceptual learning of double-vowel mixtures in musicians and nonmusicians, we found: (i) stronger alpha-band power preceding incorrect responses, especially for musicians, (ii) greater learning-related changes in connectivity for musicians, especially in the right hemisphere and preceding incorrect responses, and (iii) stronger bottom-up (auditory-to-motor) than top-down (motor-to-auditory) connectivity for both groups.

### Musicians displayed stronger modulation of alpha activity that varied with trial success

Musicians displayed stronger alpha-band (7-12 Hz) activity preceding incorrect trials in our double-vowel identification task than nonmusicians. As induced entrainment here (∼9 Hz) overlaps with the alpha range, increased alpha activity could indicate stronger persistent stimulus entrainment after the sound has stopped (exogenous activity) or *reduced* outward attention to the stimulus and greater inward reflective processing (endogenous activity) (Klimesch, 2012). Given that increased alpha-band power was observed prior to incorrect responses, the latter interpretation is more plausible. One potential explanation for increased activity associated with reduced attention to task may be that increased alpha power reflects decreased neuronal excitability (e.g., inhibition), which leads to reduced stimulus encoding at the sensory level. As a result of the stimulus not being encoded properly, task performance becomes poorer (Iemi et al., 2022). Increases in alpha activity in the pre-stimulus period could indicate that participants were “tuning out” the trial and therefore responded incorrectly (Klimesch, 2012).

Trial-dependent changes in alpha power were stronger for musicians than nonmusicians preceding incorrect trials, despite similar levels of activity between groups preceding correct responses. One explanation for this finding could be that musicians are greater “modulators” of alpha activity. Previous studies have shown that changes in alpha activity are associated with improved task performance (Klimesch, 2012; Lai et al., 2022; Pfurtscheller & Lopes da Silva, 1999; Price et al., 2019; Strauß et al., 2015). We have also recently demonstrated listeners who show less stimulus-related changes in their alpha (i.e., “low alpha modulators”) achieve poorer performance on speech-in-noise perception tasks (Price et al., 2019). Alpha *desynchronization* in sensory, task-relevant brain areas may even be paired with alpha *synchronization* over task-irrelevant areas where inhibition is necessary (Mazaheri et al., 2014). Alpha power is also associated with attentional biasing during auditory processing, including tasks involving the perception of difficult and ambiguous speech (Alain et al., 2023). Greater alpha activity preceding incorrect trials may thus reflect changes in task-related inhibition and/or attentional gating. Indeed, broad increases in prestimulus neural activation predict speech recognition errors (Vaden et al., 2015; Vaden et al., 2022). And consistent with our electrophysiological data, alpha power can be stronger preceding incorrect responses (Samaha et al., 2020). Regardless of which interpretation accounts for the alpha effects observed here, it is clear musicians recruit greater changes in alpha power between successful and unsuccessful trials (**Fig. 4b**). Given musicians’ faster performance in double-vowel identification (MacLean et al., 2024), it would appear that a more dynamic alpha range in brain activity is advantageous. This could reflect musicians’ greater flexibility in deploying attentional resources during speech perception (Strait & Kraus, 2011). This notion converges with previous findings showing that acoustic-phonetic properties of speech indexed by alpha rhythms are amplified in musicians and support more robust categorization in speech perception tasks (Bidelman, 2017).

Relatedly, under the attentional interpretation of alpha, increased alpha activity in musicians may be the result of greater “tuning-out,” or reduced attentional gating to the task during incorrect trials. Musicians’ greater alpha desynchronization for successful trials could reflect stronger attentional modulation or even greater resources for redirecting attention when needed (or desired). Indeed, both groups showed increased behavioral performance with block, but it is conceivable that performance may have become more automatized in musicians during the time course of learning. This is supported by our functional connectivity data. Musicians showed increased alpha-band auditory-motor connectivity with training block, whereas nonmusicians did not. In this vein, prior studies have also shown greater selective auditory attention for musicians in concurrent speech or “cocktail party” scenarios (Brown & Bidelman, 2023; Clayton et al., 2016; Strait et al., 2010), but see (Baumann et al., 2008), and there is also evidence that inhibitory attentional control is stronger and more efficient in musically trained individuals (Medina & Barraza, 2019).

Further support for interpretation of increased alpha power as reduced attention to task is supported by findings relating alpha power to creativity (Stevens & Zabelina, 2019). Similarly, greater auditory-motor connectivity for musicians preceding incorrect trials may be the result of increased internal reflection. Reflective processing coincides with decreased arousal/attention observed through alpha desynchronization (Klimesch, 2012). Internal reflections or “daydreaming” prior to incorrect trials could indicate more widespread, inefficient processing unrelated to the task (Fink & Benedek, 2014). Previous studies link stronger alpha activity and resting-state functional connectivity with creativity in long-term trait (Bazanova & Aftanas, 2008; Beaty et al., 2014) and short-term task-related (Stevens & Zabelina, 2019) contexts. Understanding increased alpha activity as reduced attention to task goes hand in hand with greater “tuning out” or internal reflection, though our task did not measure this phenomenon explicitly.

Determining the facilitatory or inhibitory role of alpha in concurrent speech listening, as well as how this role may be modulated by long-term music training, could inform future interventions to improve everyday complex listening skills (Gray et al., 2022). For example, the overall power and ability to modulate alpha activity to suppress irrelevant information declines in older listeners, which may render pre-target entrainment weaker and less viable as a mechanism for attentional gating (Klimesch, 1999; Vaden et al., 2012; Wöstmann et al., 2015). As implied by prior behavioral and neuroimaging studies, music engagement might help offset these age-related declines in auditory processing and help fortify the sensory-attentional mechanisms necessary for parsing complex speech mixtures (Bidelman & Alain, 2015; Lu et al., 2022; Zendel & Alain, 2009, 2012; Zendel et al., 2019).

Our study only examined induced brain-to-speech alpha entrainment following a cueing rhythmic speech train. Further exploration of the role of such induced (endogenous) alpha entrainment, both to external speech and between brain areas, and how it interacts with stimulus-related speech phase-locking (Puschmann et al., 2018) is needed in order to understand neuroplastic changes in neural entrainment and how it benefits concurrent speech perception. In this vein, neurostimulation studies have already demonstrated that enhancing cortical entrainment causally improves comprehension including performance for noise-degraded speech (Guilleminot & Reichenbach, 2022; Wilsch et al., 2018).

### Auditory-motor connectivity during concurrent speech listening differs based on long-term music training

During our active double-vowel perception task, musicians showed greater auditory-motor connectivity in the right hemisphere, whereas nonmusicians displayed similar connectivity in both hemispheres. These results are in line with emerging findings suggesting long-term music training is associated with stronger functional connectivity in the right hemisphere that is associated with preserved speech-in-noise capabilities with age (Zhang et al., 2024). Right hemispheric brain pathways are dominant for pitch and fine spectral processing (Zatorre et al., 2002; Zatorre et al., 1992). Thus, greater RH engagement in musicians may indicate their greater “cue-weighting” of pitch-based cues to distinguish vowels during our concurrent speech task, in line with our previous ERP findings of the same data (MacLean et al., 2024). Relatedly, other studies suggest that musicians have stronger right hemisphere entrainment to speech within the alpha band (Puschmann et al., 2021). Nonmusicians’ similar patterns of connectivity between left and right hemispheres may indicate that neither a left-biased linguistic (Hickok & Poeppel, 2007; Mankel et al., 2022) nor right-biased pitch strategy was preferred. Given musicians’ greater speed in the task (MacLean et al., 2024), a pitch-based, spectral strategy may have been advantageous which could explain the larger recruitment of RH activity observed in our data.

### Auditory-motor connectivity is stronger in the bottom-up vs. top-down direction

We found both musicians and nonmusicians had stronger connectivity in the auditory-to-motor compared to motor-to-auditory direction prior to double-vowel identification. The directionality of connectivity provides insight as to whether concurrent speech stimuli were processed in a bottom-up (auditory-to-motor) or top-down (motor-to-auditory) manner. Here, greater auditory-motor connectivity preceding behavior may indicate more reliance on the extraction of stimulus-specific features than anticipatory motor representations of the speech stimuli (Morillon & Baillet, 2017; Tian & Poeppel, 2012). One idea is that the motor system becomes involved in speech perception when listening becomes difficult, such as when acoustic input is sparse (Osnes et al., 2011) or speech is presented in noise (Du et al., 2016). As we observed stronger bottom-up connectivity for both groups, stimulus-based feature extraction may be more advantageous than anticipatory timing during our task. That is, the repetitive stimulus train and simultaneous onset for both vowels may have decreased the need for reliance on top-down anticipatory motor-system strategies (Wu et al., 2014). Alternatively, if the task became more automatic with learning, this would tend to evoke more bottom-up signaling, which is also enhanced in musicians (Bidelman & Krishnan, 2010; Bidelman et al., 2014; Musacchia et al., 2007; Parbery-Clark et al., 2009; Puschmann et al., 2018). Our stimulus train was also periodic and predictable. It is possible that changes to the timing and/or predictability of speech sounds (e.g., jittered stimulus train) may differentially recruit auditory-motor engagement and alter the direction of connectivity during speech processing (cf. Momtaz & Bidelman, 2024; Morillon & Baillet, 2017). Future studies are needed to test these possibilities.

## AUTHOR CONTRIBUTIONS

Jessica MacLean (Design, Data Collection, Data Analysis, Writing), Jack Stirn (Data Collection, Writing), and Gavin Bidelman (Design, Data Collection, Data Analysis, Writing).

